# Predicting synergism of cancer drug combinations using NCI-ALMANAC data

**DOI:** 10.1101/504076

**Authors:** Pavel Sidorov, Stefan Naulaerts, Jérémy Ariey-Bonnet, Eddy Pasquier, Pedro J. Ballester

**Author notes:** Correspondence to: Pedro Ballester.

## Abstract

**Background:** Drug combinations are of great interest for cancer treatment. Unfortunately, the discovery of synergistic combinations by purely experimental means is only feasible on small sets of drugs. *In silico* modeling methods can substantially widen this search by providing tools able to predict which of all possible combinations in a large compound library are synergistic. Here we investigate to which extent drug combination synergy can be predicted by exploiting the largest available dataset to date (NCI-ALMANAC, with over 290,000 synergy determinations).

**Methods:** Each cell line is modeled using primarily two machine learning techniques, Random Forest (RF) and Extreme Gradient Boosting (XGBoost), on the datasets provided by NCI-ALMANAC. This large-scale predictive modeling study comprises more than 5000 pair-wise drug combinations, 60 cell lines, 4 types of models and 5 types of chemical features. The application of a powerful, yet uncommonly used, RF-specific technique for reliability prediction is also investigated.

**Results:** The evaluation of these models shows that it is possible to predict the synergy of unseen drug combinations with high accuracy (Pearson correlations between 0.43 and 0.86 depending on the considered cell line, with XGBoost providing slightly better predictions than RF). We have also found that restricting to the most reliable synergy predictions results in at least two-fold error decrease with respect to employing the best learning algorithm without any reliability estimation. Alkylating agents, tyrosine kinase inhibitors and topoisomerase inhibitors are the drugs whose synergy with other partner drugs are better predicted by the models.

**Conclusions:** Despite its leading size, NCI-ALMANAC comprises an extremely small part of all conceivable combinations. Given their accuracy and reliability estimation, the developed models should drastically reduce the number of required *in vitro* tests by predicting *in silico* which of the considered combinations are likely to be synergistic.

## Introduction

Drug combinations are a well-established form of cancer treatment^1^. Administering more than one drug can provide many benefits: higher efficacy, lower toxicity and at least delayed onset of acquired drug resistance^2–4^. Serendipitous discovery in the clinic has been a traditional source of effective drug combinations^5,6^. Yet systematic large-scale efforts to identify them have only recently been pursued, with a growing number of preclinical experimental efforts to identify synergistic combinations^6–11^ being reported in literature. The sheer number of available and possible drug-like molecules^12^ and an exponential number of their combinations, however, make the process of finding new therapeutic combinations by purely experimental means highly inefficient.

An efficient way of discovering molecules with previously unknown activity on a given target is using *in silico* prediction methods. Quantitative Structure-Activity Relationship (QSAR) models establish a mathematical relationship between the chemical structure of a molecule, encoded as a set of structural and/or physico-chemical features (descriptors), and its biological activity on a target. Such methods have been successfully used in a wide variety of pharmacology and drug design projects^13^, including cancer research^14–16^. QSAR models are traditionally built using simple linear models^17–20^ to predict the activity of individual molecules against a molecular target. In the last 15 years, non-linear machine learning methods, such as Neural Network (NN)^21^, Support Vector Machine (SVM)^22^ or Random Forest (RF)^23^, have also been employed to build QSAR models. More recently, QSAR modeling has also achieved accurate prediction of compound activity on non-molecular targets such as cancer cell lines^24^.

To extend QSAR modeling beyond individual molecules, the set of features from each molecule in the combination must be integrated. Various ways exist to encode two or more molecules as a feature vector (*e.g.* SIRMS descriptors^25^ for properties of combinations or the CGR approach for chemical reactions^26^), and rigorous validation strategies^27^ for the resulting models have been developed. The most common representation of a drug pair is, however, the concatenation of features from both molecules^28^. On the other hand, modeling drug combinations requires the quantification of their synergy. Several metrics exist to quantify synergy^29^ (*e.g.* Bliss independence^30^, Loewe additivity^31^, Highest single agent approach^32^ or Chou-Talalay Method^33^). These are implemented in various commercial and publicly available software kits for the analysis of combination data, *e.g.* Combenefit^34^, CompuSyn (http://www.combosyn.com) or CalcuSyn (http://www.biosoft.com/w/calcusyn.htm).

One major roadblock in drug synergy modeling has been the lack of homogeneous data (*i.e.* datasets generated with the same assay, experimental conditions and synergy quantification). This has been, however, alleviated by the recent availability of large datasets from High-Throughput Screening (HTS) of drug combinations on cancer cell lines. For instance, Merck has released an HTS synergy dataset^35^, covering combinations of 38 drugs and their activity against 39 cancer cell lines (more than 20,000 measured synergies). This dataset has been used to build predictive regression and classification models using multiple machine learning methods^36^. AstraZeneca carried out a screening study, spanning 910 drug combinations over 85 cancer cell lines (over 11,000 measured synergy scores), which was subsequently used for a DREAM challenge^37,38^. Very recently, the largest publicly available cancer drug combination dataset has been provided by the US National Cancer Institute (NCI). This NCI-ALMANAC^39^ tested over 5,000 combinations of 104 investigational and approved drugs, with activities measured against 60 cancer cell lines (more than 290,000 measured synergies).

Here we take the opportunity provided by NCI-ALMANAC datasets to investigate to which extent we can predict the synergy of unseen drug combinations on NCI-60 cell lines. We build an individual model for each cell line using the popular Random Forest (RF) algorithm^40^. We also build a second model per cell line using XGBoost (XGB for short)^41^, a recent machine learning method that has helped to win numerous Kaggle competitions^41^ as well as to generate highly predictive QSAR models^42^. We validate these models for commonly-encountered prediction scenarios: *e.g.* unseen drug combination or unseen drug partner. Importantly, we also investigate the use of reliability estimation techniques to further improve prediction of drug combination synergy. Lastly, we assess the suitability of NCI-ALMANAC datasets for predictive modeling depending on the screening center where they were generated.

## Methods

### Data

NCI-ALMANAC is the largest-to-date phenotypic drug combination HTS. It contains the synergy measurements of pairwise combinations of 104 FDA approved drugs on the 60 cancer cell lines forming the NCI-60 panel^43^. The drugs include a wide array of small organic compound families, as well as several inorganic molecules (cisplatin and related platinum-organic compounds, arsenic trioxide). A similarity clustering dendrogram (Figure 1) indicates the high diversity of the drugs. Indeed, only 3 clusters comprising 8 drugs are formed with a Tanimoto score threshold of 0.8 (Vinblastine with Vincristine, Sirolimus and Everolimus, and Daunorubicin-Doxorubicin-Idarubicin-Epirubicin clusters), while the remaining 96 drugs have smaller similarity among them.

**Figure 1.**
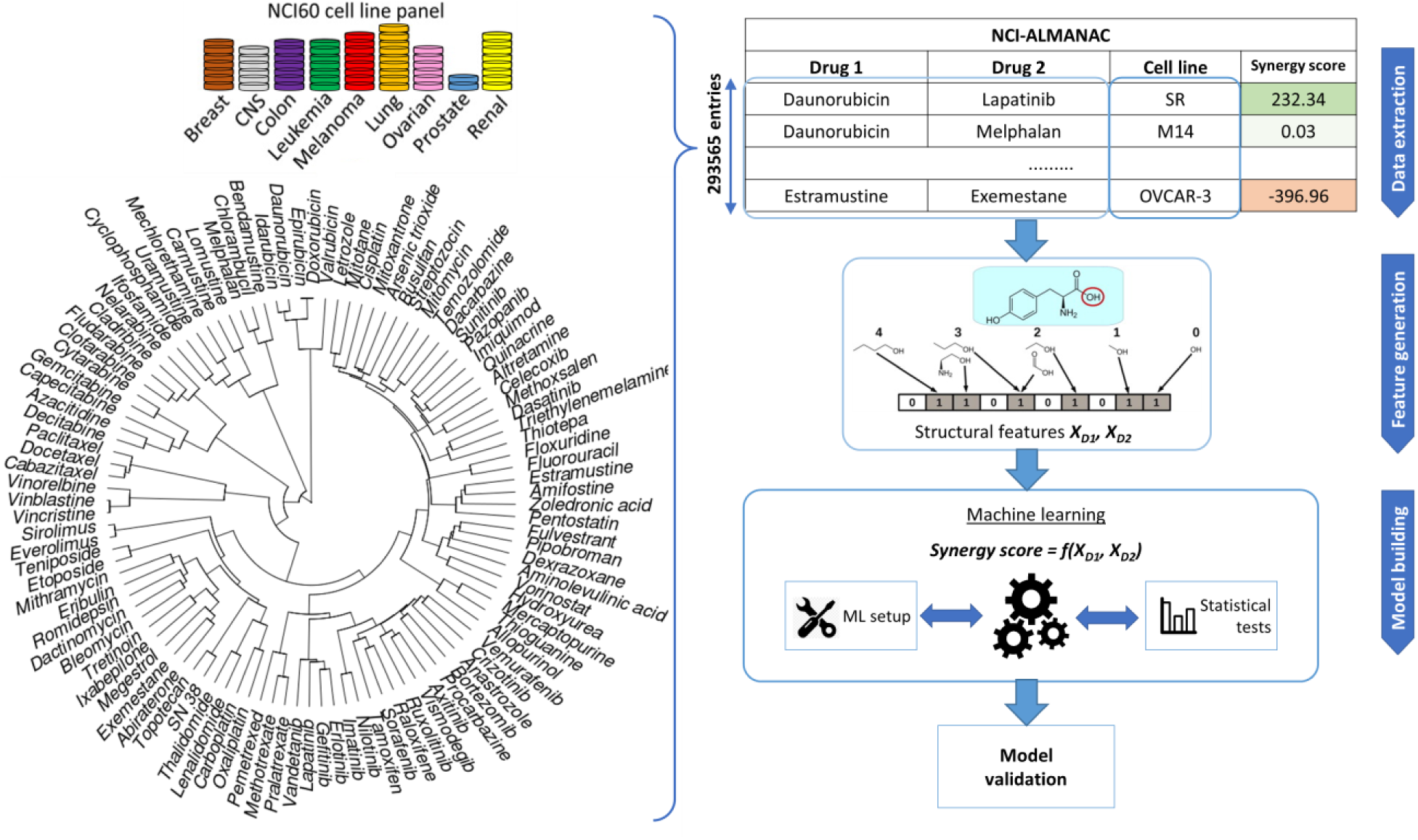
Sketch of the workflow for drug combination modeling. Training data comes from NCI-ALMANAC, which comprises over 290,000 synergy measurements from pairs of 104 drugs tested on the 60 cell lines. Similarity clustering diagram for 104 NCI-ALMANAC drugs is on the left. Each drug is characterized by Morgan fingerprints complemented with physico-chemical properties, using the Tanimoto score on these features as the similarity metric. Hierarchical agglomerative clustering was carried out (ward.D2 algorithm in the R hclust function). Closely related compounds form tight clusters (*e.g.* doxorubicin and its analogs, analogs of paclitaxel, etc). By contrast, naturally inorganic compounds such as cisplatin and arsenic trioxide appear as outliers (the highest similarity coefficient to other drugs being 0.156 and 0.125, respectively). The calculated drug features are also utilized to build and test predictive models by machine learning approach. The predictive accuracies of the models are determined by multiple cross-validation experiments.

NCI-ALMANAC aggregates synergy data from three screening centers: NCI’s Frederick National laboratory for Cancer Research (screening center code 1A, 11,259 synergy determinations), SRI International (FF, 146,147 determinations), and University of Pittsburgh (FG, 136,129 determinations). The synergy of drug pairs is measured in these screening centers against the NCI-60 panel, which includes cell lines from nine cancer types: leukemia, melanoma, non-small-cell lung, colon, central nervous system (CNS), ovarian, renal, prostate, and breast. In total, synergy is measured for 293,565 binomials drug combination – cell line, which represents a matrix completeness of 91.35%. Each center follows its own protocol, and some drugs are absent from the combination pool depending on the screening center. Since there is no overlap between drug combination – cell line pairs between the three centers, it is not possible to estimate inter-center batch effects, and therefore we must use data from different screening centers separately.

The combination benefit is quantified in NCI-ALMANAC by the so-called ComboScore (a modified version of the Bliss independence model, see Supplementary Information for details). From the entire dose-response matrix of the considered drug combination and cell line tuple, the gain (or loss) of the effect achieved by the combination over the theoretically expected value if the effect was additive is calculated. The positive values of ComboScore indicate a synergistic effect of the combination, the negative - an antagonistic effect.

### Machine learning workflow

Models are built using two machine learning algorithms: Random Forest (RF)^44^ and Extreme Gradient Boosting (XGBoost; XGB for short)^42^. Both algorithms are here based on regression trees. RF is one of the most widely used methods in QSAR modeling and has several advantages: it is relatively easy to set up, it does not require extensive hyperparameter tuning and its performance is competitive with methods that require such tuning^42^. Here we use the RF implementation from the scikit-learn Python library^45^. XGB employs the boosting learning paradigm and is often faster to train than RF. In this work, we used the publicly available Python implementation^41^ (http://xgboost.readthedocs.io/en/latest/python/). XGB is one of the most popular methods for Kaggle machine learning competitions, and is performing on par with other sophisticated methods such as Deep NNs in QSAR applications, while requiring less setup and hyperparameter tuning as well as being more efficient^42^.

The modeling workflow is sketched in Figure 1. Structural features are generated for each drug, drug pairs are represented by concatenated vectors of features. Five types of features are used: Molecular ACCess System (MACCS) keys^46^, Morgan fingerprints^47^, ISIDA fragments^48^, SIRMS^25^, with or without addition of physico-chemical properties (see supplementary information for their description). In theory, the order of the drugs in the concatenated vector should not influence the result. In practice, however, this may introduce a bias into the training set. In order to overcome such bias in drug positioning, a special data augmentation scheme suggested by Preuer et al.^36^ is considered: each row of the training set is duplicated by swapping the order of the two feature vectors associated with a drug pair.

Exploratory modeling to optimize the machine learning setup is first carried out on a single random data partition per cell line. The partition is made using scikit-learn library^45^ function, *train_test_split*. While these partitions are done for each cell line, they are preserved between experiments by fixing the random seed to allow a direct comparison of models’ performance. We also perform experiments with different test set sizes (5%, 10%, or 20% of data), leading to additional data partitions, as well as data preprocessing strategies. For each of these partitions, models are trained on the training set and evaluated on the test set.

The performance of the RF algorithm generally improves marginally with hyperparameter tuning^49^, thus we only evaluate five values for the number of trees (100, 250, 500, 750 and 1000 trees) and two values for the number of features considered in splitting a tree node (all and one-third of features). On the other hand, we employ XGB with a recommended set values for its hyperparameters (number of trees 700, maximum tree depth 6, learning rate 0.05, regularization coefficient 0)^42^. Furthermore, we also tune XGB for each cell line model by performing a 5-fold CV on the training set with a grid search of these four hyperparameters: number of trees (200 to 1000), maximum tree depth (5 to 10), learning rate (0.1, 0.05, 0.01) and regularization coefficient (0, 0.0001, 0.001, 0.01). The optimal values are used to build the XGB model with the entire training set of that cell line.

To evaluate a model’s performance, the following metrics are calculated from observed and predicted ComboScore values: Root Mean Squared Error (*RMSE*), Coefficient of determination (*R*^2^), Pearson’s correlation coefficient (*R*_*p*_) and Spearman’s rank-order correlation coefficient (*R*_*s*_). These metrics are always calculated on a test set not used to train or select the model. The supplementary information provides the expressions of these common metrics.

## Results

### Exploratory modeling of NCI-ALMANAC data

First, we perform an exploratory modeling on the FG datasets in order to determine optimal settings for synergy prediction by assessing various types of features, data augmentation schemes and machine learning methods. The summary of performance improvements is shown on Figure 2. The best median R_p_ across cell lines for RF was obtained with 250 trees, a third of the features evaluated at each tree node, training data augmentation and MFPC fingerprints complemented by physico-chemical properties (256 and 7 features per drug, respectively). The gain of performance with RF is substantial: the median R_p_ increases from 0.530 (I) to 0.634 (VI).

**Figure 2.**
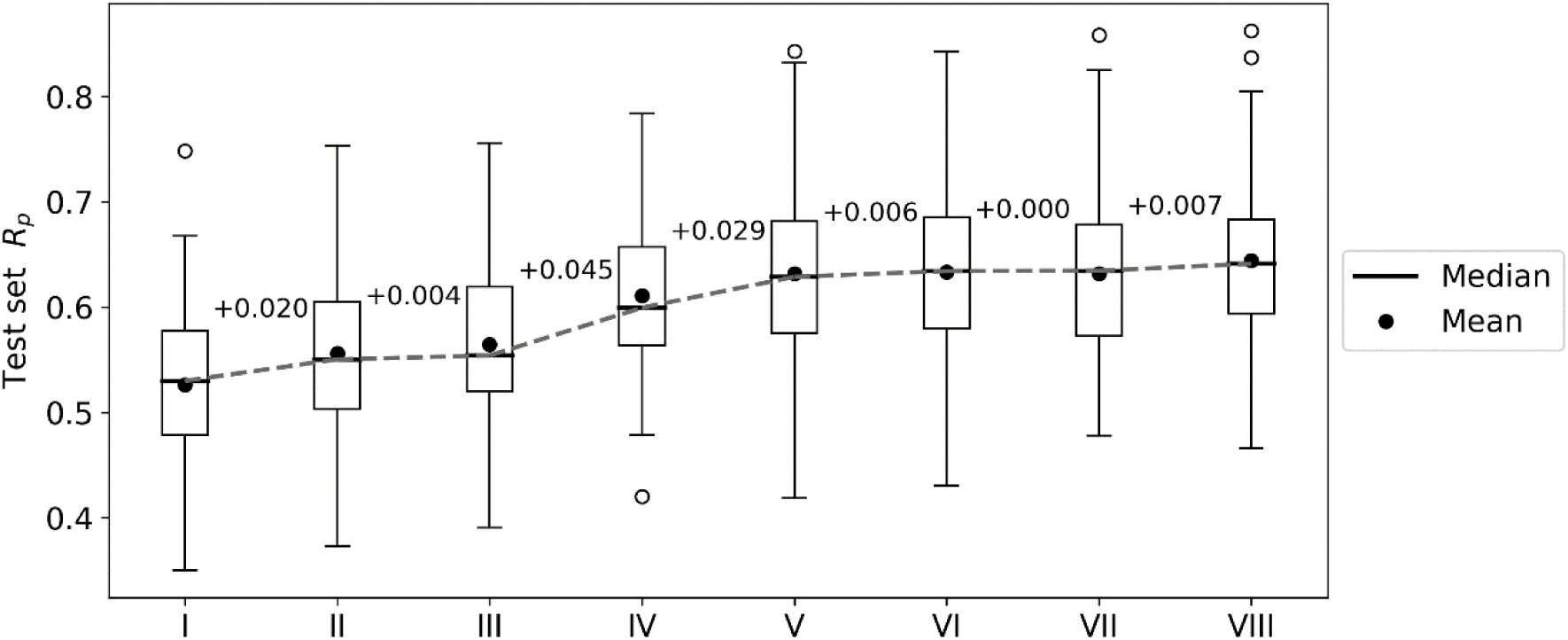
Performance gain across cell lines for each introduced modeling choice during the exploratory analysis of FG data. Each boxplot represents the distribution of the cell line models’ test set performances (R_p_) at any given step. Analysis steps are carried out sequentially: I – RF, 1000 trees with all n features tried to split a node, 80% training set, 20% test set, MACCS (Molecular ACCess System) keys as features; II – MFPC (Morgan fingerprint counts) are used as features instead; III – physico-chemical features are added for each drug; IV – training set rows are duplicated with the reverse order of drugs (data augmentation); V – 90% training set, 10% test set are used instead of the initial 80/20 partition; VI – RF with 250 trees with n/3 features tried to split a node; VII – XGB models with recommended settings; VIII – tuned XGB models. Note that I-V employ RF with same values for its hyperparameters (RF tuned in VI) and V-VIII use the same training and test sets. Modeling choices introducing the largest improvements are the choice of molecular features and the data augmentation strategies.

XGB models are generated with the same features and data set partitions. Changing the machine learning algorithm from RF to XGB does not improve the median test set R_p_, although both minimum and maximum R_p_ are higher with XGB (boxplots VI and VII in Figure 2, respectively). After tuning of XGB hyperparameters per cell line, a small gain in overall performance is obtained: the median R_p_ of tuned XGB rises to 0.641 (boxplot VIII). In comparison, Y-randomization^50^ tests using the same learning algorithm did only obtain a median R_p_ of −0.016 (−0.024 when using RF). Figure 3 shows the degree of accuracy achieved by each algorithm for the best and the worst predicted cell line. The cell lines with the worst predictions (OVCAR-8 for RF and SF295 for XGB) have substantially smaller variance in observed ComboScore than those with the best predictions (SK-MEL-5 for both algorithms).

**Figure 3.**
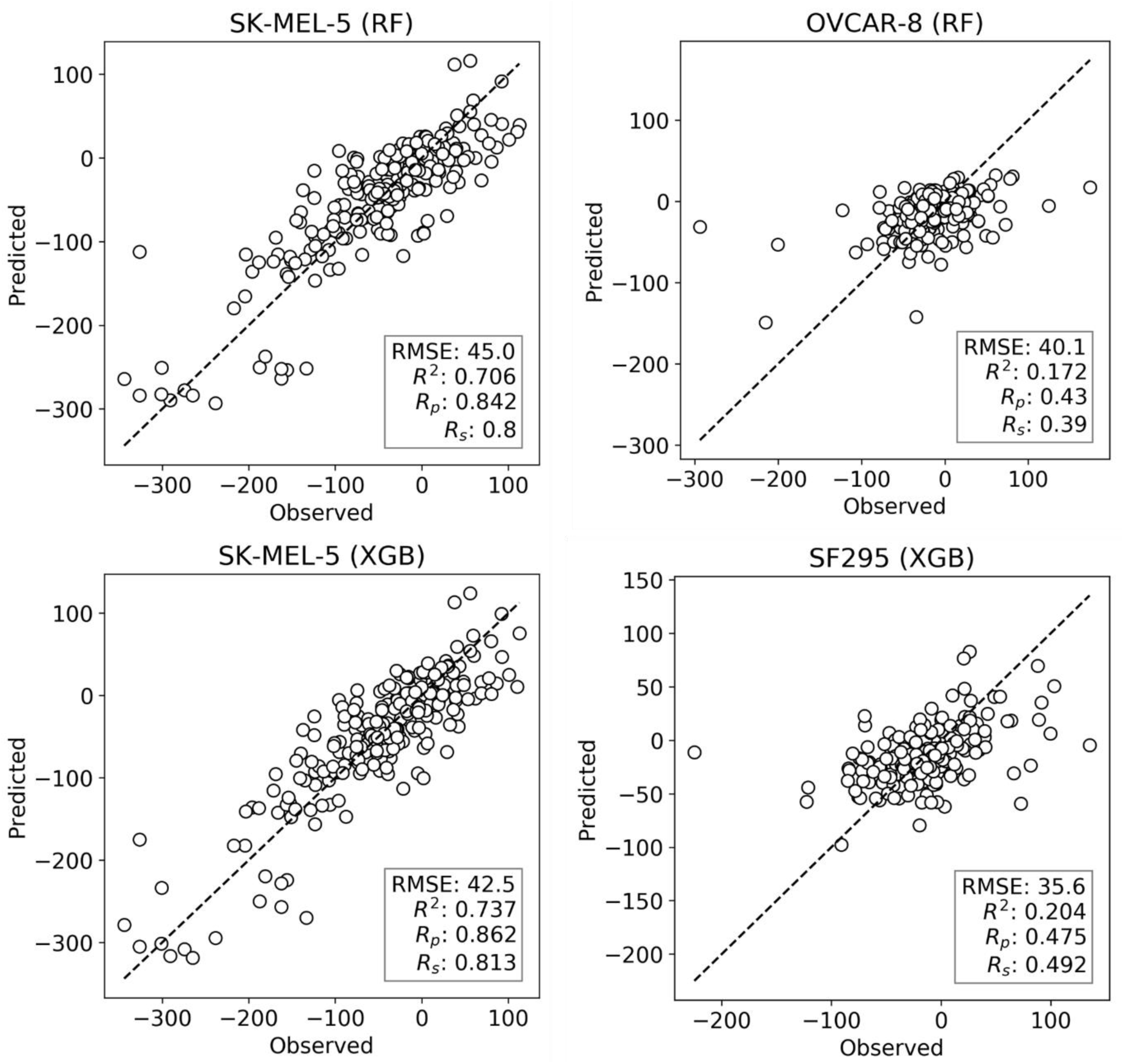
Observed vs predicted ComboScore for all the drug combinations in the test set. This is presented for the best- and worst-performing models with both ML methods, RF and XGB (these models correspond hence to the extremes of Figure 2’s boxplots VI and VIII, respectively). On the left column, the best-performing cell line models from each method. On the right side, the worst-performing cell line models. All performance metrics are shown. Each point represents a drug combination in that test set.

### Estimating the reliability of drug synergy predictions

For prospective use of models, it is paramount to calculate not only predicted drug combination synergies, but also how reliable these predictions are^51^. With this purpose, we have applied a RF-specific reliability prediction approach, where the degree of agreement between the diverse trees in the forest serves as a reliability score. This is quantified here as the standard deviation (SD) of the RF tree predictions (250 per drug combination and cell line) and referred to as tree_SD. tree_SD has been pointed out as one of the most powerful metrics to assess the reliability of predictions in regression problems^51^. We thus assemble test subsets with the 25% most reliable ComboScore predictions per cell line (*i.e.* combinations with the 25% lowest tree_SD scores). Likewise, we assemble test subsets with 25% least reliable predictions per cell line.

Figure 4 presents the test set performances of each cell line model on the three scenarios: 25% most reliable predictions, all predictions regardless of estimated reliability and 25% least reliable predictions. The top and bottom 25% predictions in terms of reliability obtain the lowest and highest RMSE in every cell line, which demonstrates the accuracy and generality of tree_SD as a reliability score for drug synergy predictions. Test set RMSE varies greatly across cell lines, *e.g.* models built on leukemia cell lines obtain in general higher error. This, however, comes from the higher range of ComboScores observed in these cell lines. Indeed, the larger this range, the higher the range of predicted ComboScores is, which combined tend to make RMSE larger. Similar RMSE is only obtained on the K-562 leukemia cell line, which is consistent with the fact that it has the lowest range among leukemia cell lines and similar to that of other cancer types.

**Figure 4.**
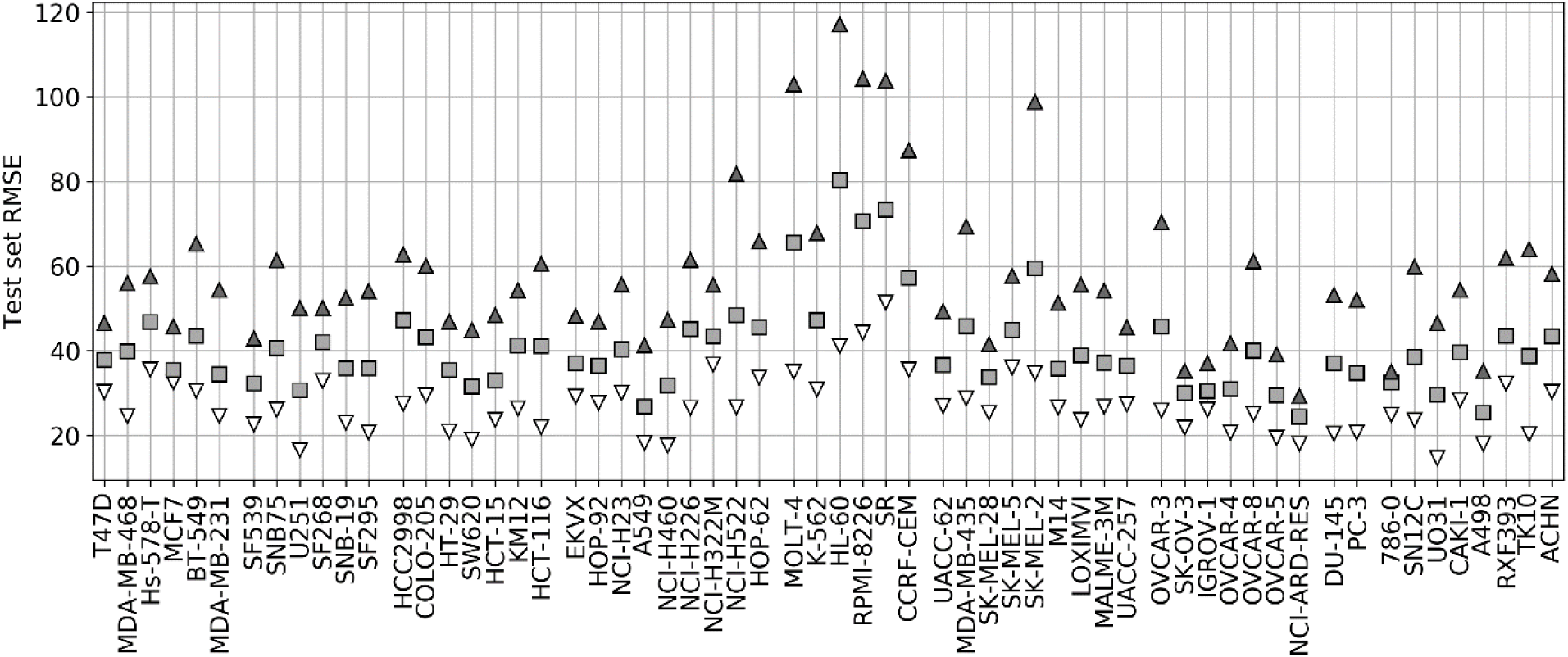
10% test set RMSE of RF cell line models trained on 90% of the FG data. Grey squares represent the model’s RMSE on all the test combinations (RF predictions as usual). Black triangles mark the RMSE of the 25% least reliable (highest tree_SD) combos, whereas white inverted triangles correspond to the RMSE of the 25% most reliable (lowest tree_SD). Cell line models in the horizontal axis are segregated by cancer type. In each cell line, the reliability score tracks test RMSE and hence it can be used to identify *a priori* the most accurate predictions.

Reliability estimation is evaluated in terms of RMSE rather than R_p_. While RMSE is not as intuitive as correlation, correlations may be misleading when comparing performances of models across test sets with distinct variances. Figure 5 illustrates this issue with the test performances of HL-60 models, which benefit the most from reliability estimation. The test set with the most reliable combinations is predicted with half the RMSE of the entire test set (RMSE of 41 vs 80) and a third of the least reliable combinations (RMSE of 41 vs 117). This more accurate prediction can be visually observed too, but the other metrics (R^2^, R_p_ and R_s_) do not capture this increase in accuracy due to substantially different ComboScore variance between the compared test sets. Importantly, RF with reliability prediction provides a much larger reduction in RMSE than that introduced by XGB (bottom right), both with respect to RF without reliability prediction (bottom left). These results strongly suggest that, in cases where it is not necessary to test all positive predictions (here synergistic drug combinations), selecting the most reliable predictions is more effective than using the most suitable ML algorithm.

**Figure 5.**
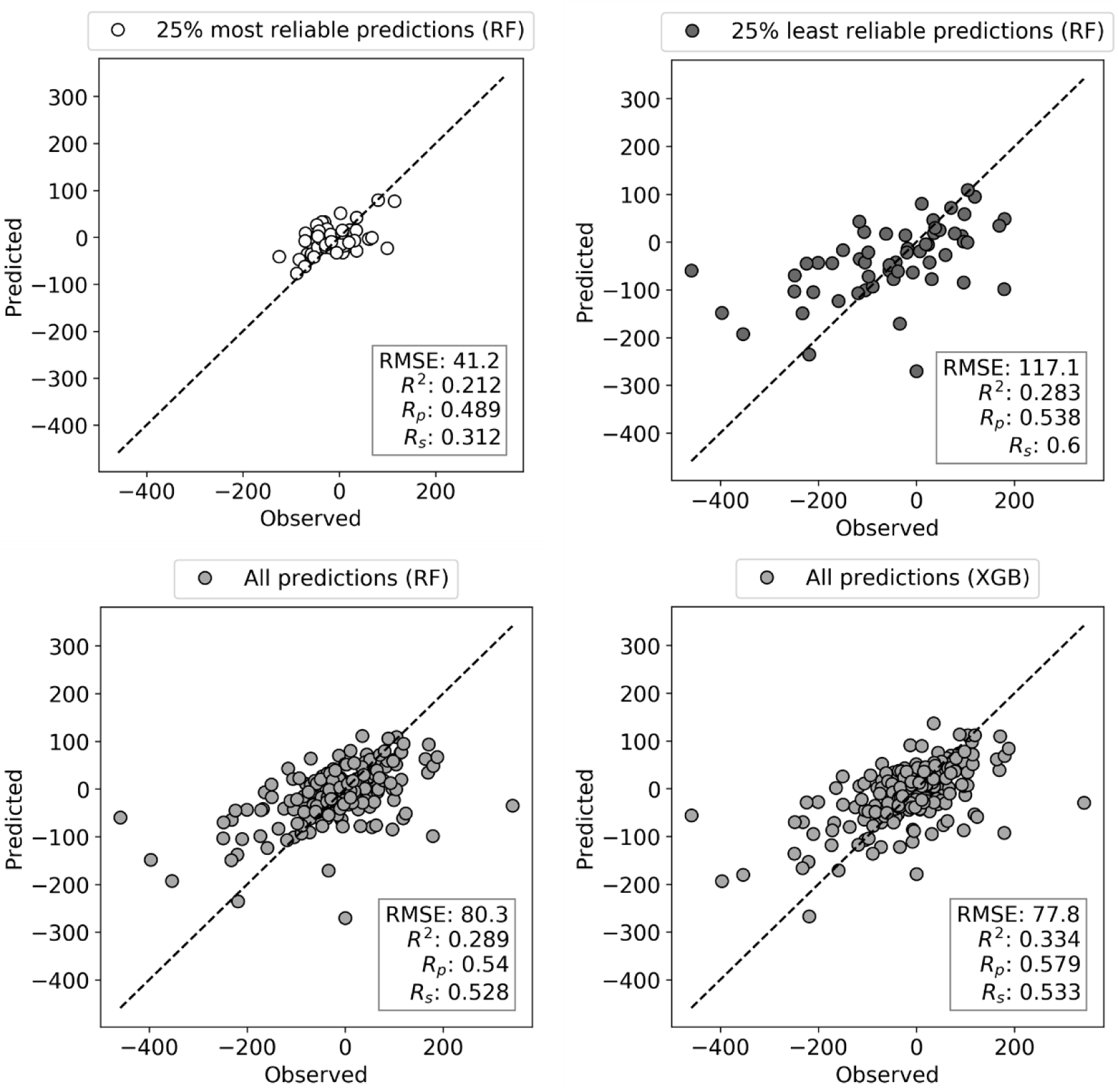
Observed vs. Predicted ComboScore plots for HL-60 leukemia cell line test set (10% of data). Models are built on 90% of FG dataset corresponding to this cell line using RF and XGB methods, both tuned. Each circle is now a drug combination from the entire test set, with its color indicating one of the three scenarios (as in Figure 4). All performance metrics are displayed in each plot. The subset with the most reliable ComboScore RF predictions (top left plot) achieves half the RMSE of the entire test set (bottom right). Importantly, this is a much larger reduction in RMSE than that introduced by XGB (bottom right) with respect to RF (bottom left). Furthermore, the most reliable predicted ComboScores (top right) obtain a third of the RMSE of the least reliable predictions (top left).

### Performance in predicting synergies with drugs not included in NCI-ALMANAC

The random data splits that we have used so far may overestimate the model’s performance in the case of drug combinations. This would be due to the presence of the two drugs in the combination in both training and test sets, albeit with other partners^27^. In order to assess to which extent this is the case, we also carry out Leave-One-Drug-Out (LODO) cross-validation experiments for each cell line. In LODO cross-validation, every combination containing a certain drug is left out as a test set, and the model is built on the remaining combinations tested on that cell line. In this way, the validation simulates the model’s behavior when presented with a new chemical entity outside of the model’s scope, as if it was not included in the dataset.

Figure 6 shows the outcome of LODO cross-validation for XGB per cell line. We henceforth use XGB with the recommended values for hyperparameters, as tuning them for each LODO cross-validation fold and cell line is prohibitive and would only provide marginal gains (see Figure 2). LODO results show that combinations associated with 75% of the left-out drugs can be predicted with an accuracy of at least R_p_ = 0.3 against any cell line. This accuracy raises to at least R_p_ = 0.5 for 50% of the left-out drugs. The latter is not much worse than the median R_p_ across cell lines when using 90/10 data partitions (R_p_ = 0.641 as shown in Figure 2’s boxplot VIII). k-fold cross-validation results are available for comparison in Supplementary Figure 3.

**Figure 6.**
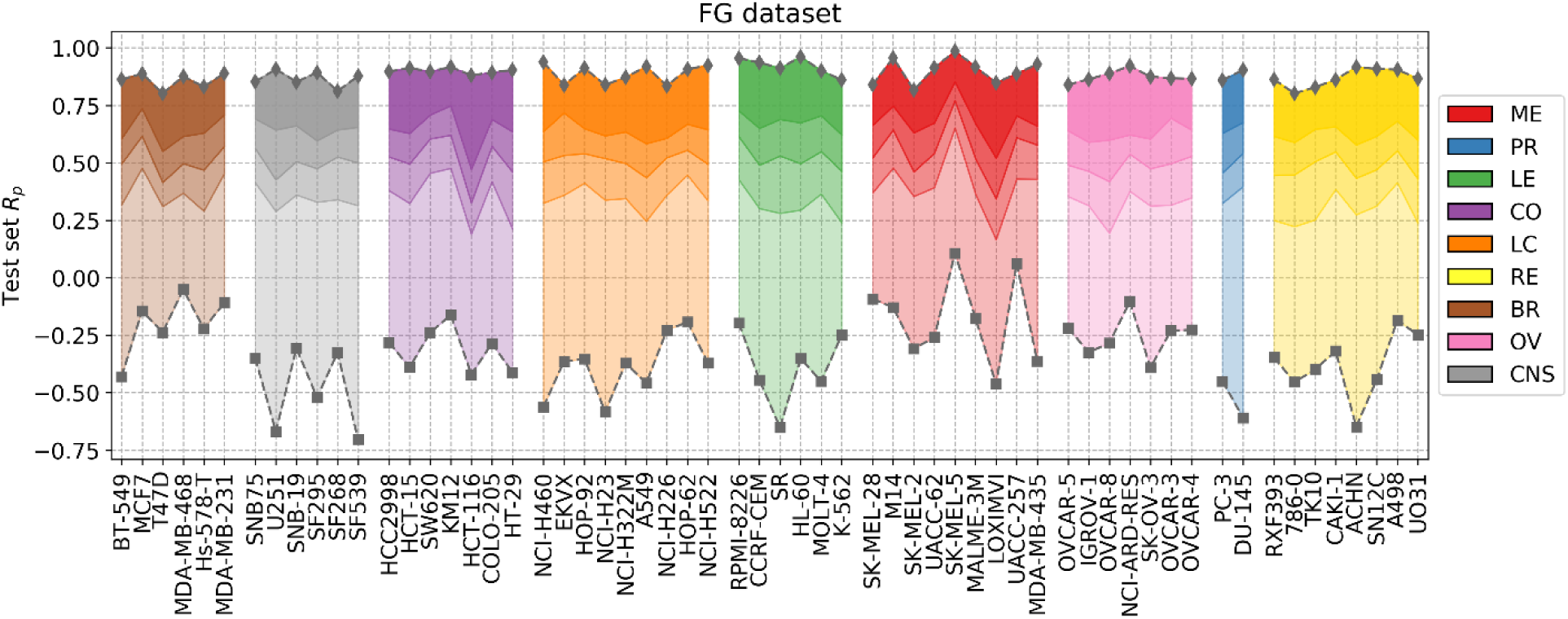
LODO cross-validation results using XGB with the recommended values for their hyperparameters on the FG dataset. Distribution of models’ performances is shown by cancer type (color code). Each colored zone represents 25% of models per cell line: from dense zone – top performing 25%; to light zone – bottom quartile. The method performs with at least moderate accuracy (R_p_>0.30) in 75% of left-out drugs (the top 3 quartiles) across cell lines. Left-out drugs within the top quartile, darkest shade among the four per cell line, are predicted with a R_p_ ranging from 0.471 (HCT-116) to 0.986 (SK-MEL-5). Although there are no large differences in how well different cancer types are predicted, left-out drugs on melanoma (ME, in red) and leukemia (LE, in green) cell lines obtain slightly higher average performance (median R_p_ of drug-out models for corresponding cell lines are 0.554 and 0.524, respectively).

However, as expected, LODO results are much worse for certain left-out drugs in each cancer type. LODO performance of each drug across cell lines is shown in Supplementary Figure 4. This figure shows that models for arsenic trioxide, highly dissimilar to other drugs, have the lowest performance across cell lines (median R_p_ of models concerning this drug is −0.28). Conversely, partners of tyrosine kinase inhibitors, well-represented in these datasets, are predicted with high accuracy (*e.g.* models for imatinib have median R_p_=0.82).

Figure 7 shows the analysis for LODO cross-validations in terms of RMSE. About 75% of models demonstrate at least moderate accuracy (RMSE<50). The exceptions are mostly leukemia cell line models, which obtain higher RMSE due to having the highest variances in ComboScores among cancer types. Importantly, using RF models restricted to the most reliable predictions allows us to reduce the error of prediction further in every cell line (RMSE<40), in full agreement with the findings from random 90/10 partitions (see Figure 4).

**Figure 7.**
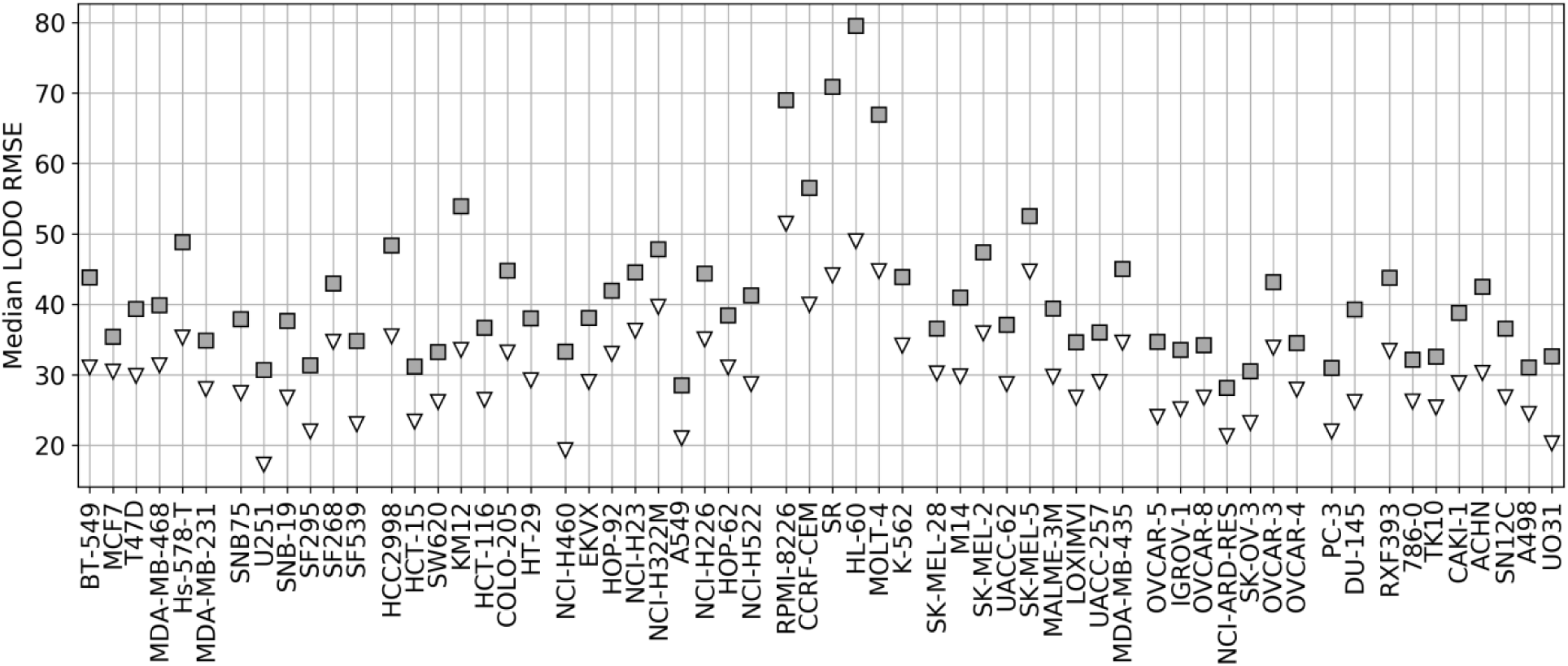
Median RMSE in LODO cross-validation for XGB with the recommended values for their hyperparameters (grey squares) and RF top 25% most reliable predictions (white inverted triangles) for each cell line (grouped by cancer type). As the plot shows, combinations with one left-out drug can be predicted with at least moderate accuracy across cell lines (RMSE < 50 for XGB, RMSE < 40 for RF with reliability estimation; both being approximate thresholds).

### Comparing predictive models built with data from different screening centers

So far we have exclusively employed data from the FG screening center, which represents about half of NCI-ALMANAC data. Practically all the remaining ComboScores come from the FF screening center and are also determined with a 3×3 concentration grid. Thus, we evaluate here the predictive potential of FF datasets. We start by building RF models from FF data using the same 90/10 partitions as with FG. Surprisingly, FF-based models obtained worse performance in every cell line (Figure 8) and thus were objectively worse at predicting ComboScores.

**Figure 8.**
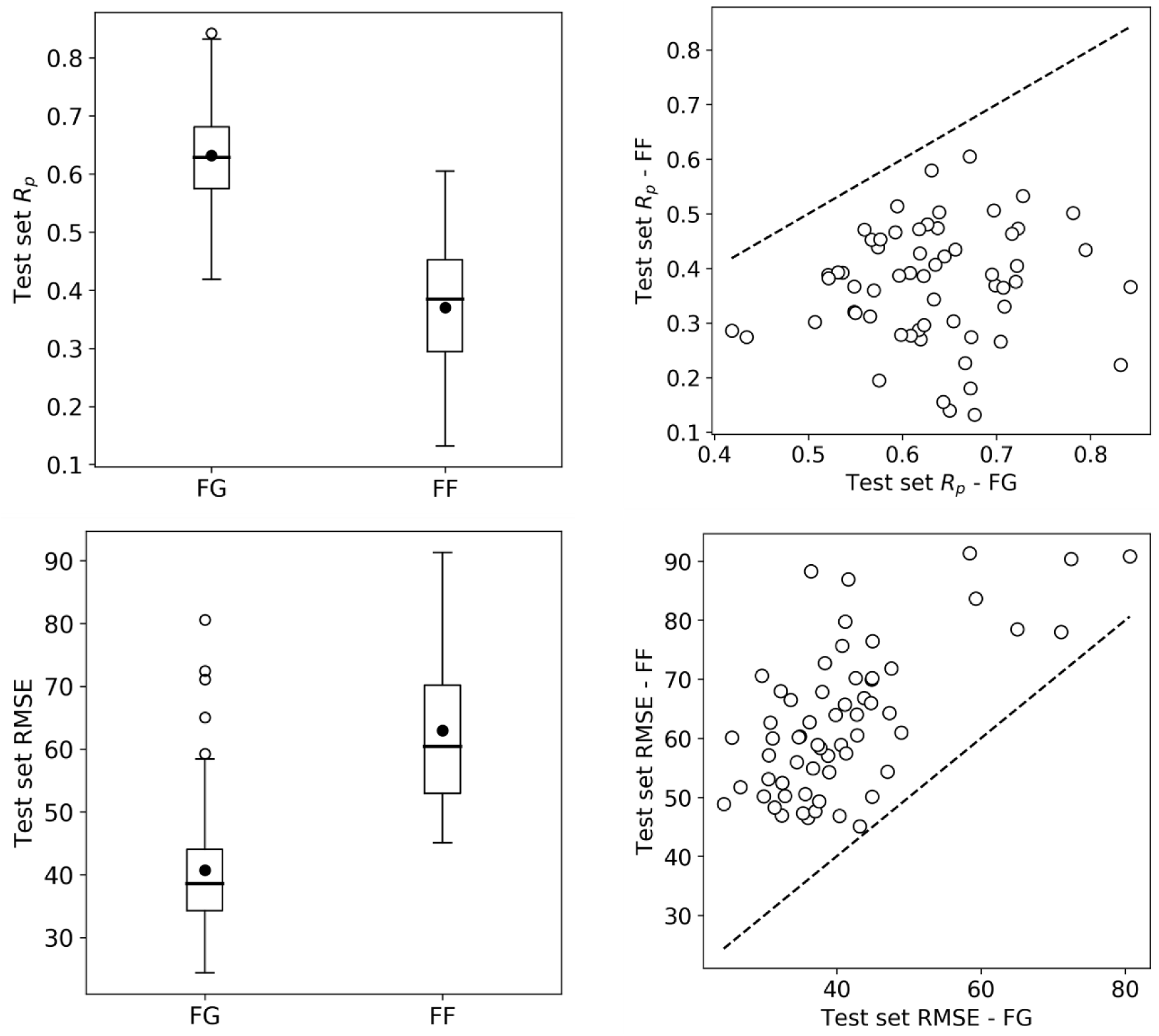
Random Forest (RF) model performance comparison for FG and FF datasets. Models are built following the final setup in the exploratory analysis (a 90/10 data partition is employed for each cell line); MFPC with physico-chemical features as well as data augmentation are also used). On the left, boxplots for cell line models test set R_p_’s (top row) and RMSE (bottom row) for both centers data. On the right, R_p_ (top row) and RMSE (bottom row) of models trained on FG dataset against models trained on FF dataset, each point shows the two model performances for the cell line. FF models obtain consistently lower performance than FG models. As the same modelling workflow was used, this strongly suggests that FF data is less predictive than FG data.

In trying to understand this surprising result, we started by investigating whether this was due to modeling differences, but this was not the case. First, FF training sets are slightly larger than FG datasets (see supplementary table 1), which theoretically favors better performance on FF. Furthermore, using tuned XGB models led to essentially the same result (median R_p_ of 0.641 for FG vs 0.368 for FF) as shown in Figure 8 with RF. In addition to these nonlinear methods, we also used Elastic Net (EN), but FF models were still substantially less predictive than FG models (median R_p_ of 0.37 for FG vs 0.23 for FF). When we carried out LODO cross-validations instead of 90/10 partitions, the same trend was observed (supplementary Figures 5 and 6 also show worse performance of FF-based LODO than that of FG-based LODO in Figure 6).

To shed light into this issue, we looked at the only factor that we can compare between these screening centers: the relative growth inhibition (PERCENTGROWTH) induced by a given concentration of a drug tested individually. Interestingly, by counting the different test dates, we observed that FG had on average tested a non-combined drug 3.77 times per cell line, whereas FF almost doubled this number (7.13 times per cell line). A higher number of tests is not in itself worrisome if the growth inhibition of the drug-concentration-cell line tuple is similar between dates. However, if the measurements from these tests are substantially different, this is a problem because the set of ComboScores determined with variable measurements from the same tuple will be inconsistent as well. Consequently, synergy differences between such combinations will not only come from their intrinsic properties, but also from unrelated experimental variability.

To show that higher growth inhibition variability in FF data results in less predictive models, we analyzed five drugs (Thioguanine, Chlorambucil, Altretamine, Fluorouracil and Melphalan) with a high number of different test dates in both centers. We first consider the drugs on a cell line were only FG models obtain high average accuracy in predicting synergy (NCI/ADR-RES) and subsequently on another were both FF and FG models are on average predictive (NCI-H322M). On each cell line, each drug has a set of growth inhibition replicates per concentration and screening center (*i.e.* 15 sets per screening center). The performance on NCI/ADR-RES using FF data is indeed poor (R_p_=0.14 in 90/10 partition by RF), but it is much better predicted using FG data (R_p_=0.65, using the same partition and method). 14 of the 15 sets have higher standard deviation of growth inhibition with FF data (Supplementary Figure 7), which is consistent with the lower accuracy in predicting synergy obtained with this dataset. Conversely, we repeated this operation with NCI-H322M where synergy is well predicted by RF with both FF (R_p_=0.61 in 90/10 partition) and FG data (R_p_=0.66, on the same partition). The standard deviations from both screening centers are now similar (Supplementary Figure 7). Taken together, these experiments suggest that the reason why FF data results in less predictive models is the noise introduced in ComboScore determination by larger variability of growth inhibition measurements.

## Discussion

NCI-ALMANAC is an extremely valuable resource for the discovery of novel synergistic drug combinations on NCI-60 cell lines. First, it is by far the largest-to-date HTS of drug combinations, therefore allowing *in silico* models with much higher accuracy and domain of applicability in predicting the synergy of other combinations. Second, some of the synergistic drug combinations discovered *in vitro* by NCI-ALMANAC were subsequently tested on human tumor mouse xenografts of the same cell line. 48% of them were also synergistic in at least one of these *in vivo* models^39^, which led to the launch of two clinical trials so far (NCT02211755 and NCT02379416).

In this study, we have found that it is possible to predict the synergy of unseen drug combinations against NCI-60 panel cell lines with high accuracy by exploiting NCI-ALMANAC data. We have established a general ML workflow (types of structural features, data preprocessing strategy, ML method) to generate such models. When trained on FG data, predicted synergies from these models match observable synergies with R_p_ correlations between 0.43 and 0.86 depending on the considered cell line. Incidentally, these regression problems must be highly nonlinear, as EN leads to substantially less predictive models than XGB or RF.

Some cell lines and drug combinations can be predicted with higher accuracy than others. For example, models for the SK-MEL-5 cell line perform best with any method (Figure 6). However, if we use RMSE instead of R_p_ to reduce the influence of the ComboScore range, models for the NCI-ARD-RES are now best (grey squares in Figure 7). LODO cross-validation also revealed both best and worst partner drugs. These differences are mainly due to the number of similar partner drugs. For example, it is difficult to predict synergy of combinations containing arsenic trioxide because its 103 partner drugs are highly dissimilar in terms of chemical structure and physico-chemical properties. Indeed, machine learning from dissimilar data instances tend to be less accurate, although here the dissimilarity can be partial as arsenic trioxide’s partner can be similar to other NCI-ALMANAC drugs. On the other hand, combinations containing some other drugs are better represented in NCI-ALMANAC and hence tend to be predicted with higher accuracy. This is the case of various alkylating agents, tyrosine kinase inhibitors and topoisomerase inhibitors (Supplementary Figure 4).

Recent QSAR and drug combination modeling studies have evaluated the application of the latest machine learning algorithms (e.g. XGBoost, Deep NN). These studies have found that these algorithms provide better performance on average across targets than RF, although these gains are small and do not always justify the much greater resources required for hyperparameter tuning^36,42^. Performance gains have also been found small here with NCI-ALMANAC data, as the average test set R_p_ of XGBoost across the 60 cell lines is just +0.007 larger than with RF. An important result is that restricting to the most reliable RF predictions provides much larger predictive accuracy than that introduced by a more suitable learning algorithm (e.g. XGBoost). It is surprising that this powerful technique is so uncommonly used, as has already been pointed out^51,53^. In fact, we are not aware of any other previous study applying reliability estimation to the prediction of drug synergy on cancer cell lines. Here reliability prediction permitted to reduce the RMSE by up to 50% depending on the cell line. This is particularly exciting for virtual screening problems, where only a small subset of the predictions can be tested *in vitro*. In this scenario, it is useful to identify those combinations that are not only predicted to be synergistic, but also reliable because this should provide higher hit rates. Lastly, highly synergistic combinations predicted with low reliability should be also tested, as the corresponding measurements will be broadening the applicability domain of future models the most.

We have also found that using FG datasets leads to substantially more predictive models than FF datasets. This result is robust in that it is observed with various types of models (XGB, RF, EN). Moreover, it occurs in spite of the availability of slightly more training data. Further investigation revealed that there are many more measurements of growth inhibition and with greater variability in FF than in FG. This in turn introduces more noise into ComboScore determinations in FF, thus impairing its modeling. Inconsistencies between centers measuring the response of drugs on cancer cell lines have been observed before^54^. There has been intense controversy about the extent, sources and impact of these inconsistencies^55–58^. In any case, it is clear that data permits the development of predictive models regardless of the screening center^59–62^, as it has also been the case here with NCI-ALMANAC. Owing to this controversy on datasets from multiple screening centers, a better understanding of their limitations and the identification of protocols to generate them with improved consistency has emerged^63^. These protocols will ultimately permit merging datasets from different screening centers to gain further predictive accuracy, which is currently not advantageous due to the discussed limitations.

While NCI-ALMANAC measured the synergies of over 5000 combinations per cell line, this still represents a minuscule part of all conceivable combinations. Even if we restricted ourselves to the set of 12,000 drug molecules estimated to have reached clinical development or undergone significant preclinical profiling^52^, almost 72 million combinations per cell line would have to be tested *in vitro* to identify the most synergistic among them. Therefore, the developed *in silico* models are of great importance because these can drastically reduce the number of required *in vitro* tests by predicting which of the considered combinations are likely to be synergistic.

