## Supplementary information for "Predicting synergism of cancer drug combinations using NCI-ALMANAC data"

### NCI-ALMANAC composition

**Supplementary Table 1.** Number of measured ComboScore values in NCI-ALMANAC dataset, grouped by cancer types, cell lines, and screening centers. In total, there are 293,565 ComboScore values across 5050 drug combinations.

| Cell line | | FG | FF | 1A |
|---|---|---|---|---|
| Breast | BT-549 | 2280 | 2480 | 183 |
| | Hs-578-T | 2256 | 2471 | 201 |
| | T-47D | 2265 | 2467 | 185 |
| | MCF7 | 2289 | 2518 | 201 |
| | MDA-MB-468 | 2274 | 2475 | 182 |
| | MDA-MB-231 | 2288 | 2513 | 197 |
| CNS | U251 | 2281 | 2498 | 199 |
| | SF295 | 2290 | 2333 | 200 |
| | SNB-19 | 2283 | 2509 | 201 |
| | SNB75 | 2282 | 2277 | 201 |
| | SF268 | 2283 | 2516 | 201 |
| | SF539 | 2274 | 2414 | 199 |
| Colorectal | SW620 | 2289 | 2533 | 201 |
| | COLO-205 | 2273 | 2450 | 199 |
| | HT-29 | 2279 | 2495 | 201 |
| | HCT-15 | 2290 | 2510 | 200 |
| | KM12 | 2247 | 2500 | 199 |
| | HCT-116 | 2285 | 2471 | 192 |
| | HCC2998 | 2226 | 2476 | 196 |
| Lung cancer | A549 | 2286 | 2474 | 194 |
| | EKVX | 2287 | 2505 | 1 |
| | HOP-62 | 2285 | 2355 | 201 |
| | NCI-H322M | 2257 | 2477 | 199 |
| | NCI-H226 | 2285 | 2488 | 192 |
| | NCI-H23 | 2283 | 2510 | 201 |
| | NCI-H460 | 2274 | 2456 | 201 |
| | HOP-92 | 2285 | 2383 | 137 |
| | NCI-H522 | 2236 | 2449 | 175 |
| Pros-tate | PC-3 | 2283 | 2497 | 185 |
| | DU-145 | 2274 | 2497 | 198 |

| Cell line | | FG | FF | 1A |
|---|---|---|---|---|
| Leukemia | CCRF-CEM | 2247 | 2420 | 190 |
| | RPMI-8226 | 2129 | 2446 | 200 |
| | K-562 | 2283 | 2434 | 173 |
| | SR | 2248 | 2402 | 190 |
| | MOLT-4 | 2198 | 2415 | 177 |
| | HL-60 | 2243 | 2234 | 165 |
| Melanoma | UACC-257 | 2284 | 2519 | 184 |
| | LOXIMVI | 2259 | 2318 | 159 |
| | MDA-MB-435 | 2226 | 2451 | 196 |
| | UACC-62 | 2281 | 2478 | 199 |
| | M14 | 2274 | 2461 | 199 |
| | SK-MEL-2 | 2188 | 915 | 167 |
| | SK-MEL-5 | 2272 | 2471 | 199 |
| | SK-MEL-28 | 2283 | 2512 | 201 |
| | MALME-3M | 2288 | 2499 | 153 |
| Ovarian | SK-OV-3 | 2278 | 2513 | 200 |
| | OVCAR-8 | 2278 | 2489 | 200 |
| | OVCAR-5 | 2289 | 2515 | 200 |
| | NCI-ADR-RES | 2289 | 2544 | 169 |
| | OVCAR-4 | 2256 | 2475 | 193 |
| | IGROV1 | 2281 | 2479 | 200 |
| | OVCAR-3 | 2260 | 2399 | 151 |
| Renal | SN12C | 2283 | 2491 | 201 |
| | RXF_393 | 2278 | 2399 | 185 |
| | A498 | 2289 | 2491 | 200 |
| | CAKI-1 | 2284 | 2463 | 185 |
| | TK-10 | 2271 | 2506 | 198 |
| | ACHN | 2282 | 2527 | 201 |
| | 786-0 | 2287 | 2462 | 201 |
| | UO-31 | 2252 | 2452 | 201 |

**NCI-ALMANAC screening centers**

1) NCI Frederick National Laboratory (screening center code 1A) uses the NCI-60 testing protocol (https://dtp.cancer.gov/discovery_development/nci-60/methodology.htm), with 5 concentrations per single agent, and 5x3 matrices for combinations. The growth percentage is measured through the classical sulforhodamine cytotoxicity assay[1], in which the amount of bound sulforhodamine is observed absorbance measurement at 510 nm wavelength in colorimetry. Therefore, the number of viable cells is proportional to optical density of the dye. 11,259 values in total. Absent drugs: Vemurafenib, Fludarabine.

2) SRI International (FF) uses the modified protocol: drugs are tested in 3 concentrations as single agents, and in a 3x3 concentration matrix for combinations. Cell viability is measured in CellTiter-Glo luminescence assay, luminescence produced is proportional to the number of viable cells. There are 146,177 measured values in total. Absent drugs: Idarubicin, Epirubicin, Eribulin, Abiraterone, Pazopanib, Vismodegib, Crizotinib, Axotinib, Vandetanib, Vemurafenib, Ruxolitinib, Cabazitaxel.

3) University of Pittsburgh (FG) also follows a modified version of the NCI-60 test protocol, with 3x3 concentration matrices for combinations. There are 136,129 measured values from this center. Absent drugs: Doxorubicin, Epirubicin, Idarubicin, Eribulin, Triethylenemelamine.

**NCI-ALMANAC ComboScore**

Expected tumor growth percentage $Z$ for cell line $i$, after two-day treatments with drugs $A$ and $B$ at concentrations $p$ and $q$, respectively, is calculated from the observed effect of these drugs as single agents in these concentrations ($Y_i^{A_p}, Y_i^{B_q}$, truncated at 100) with the following formula:

$$Z_i^{A_pB_q} = \begin{cases} \min\left(Y_i^{A_p}, Y_i^{B_q}\right), if\ Y_i^{A_p} \leq 0\ or\ Y_i^{B_q} \leq 0 \\ \frac{1}{100}\left(Y_i^{A_p} \times Y_i^{B_q}\right), otherwise \end{cases}$$

The final ComboScore ($CS$) for the cell line and the combination is calculated as the sum of the differences between expected ($Z_i^{A_pB_q}$) and observed ($Y_i^{A_pB_q}$) effects of drug combinations at each concentration:

$$CS_i^{AB} = \sum_{p,q}\left(Z_i^{A_pB_q} - Y_i^{A_pB_q}\right)$$

Since the observed value is the tumor growth percentage, the lower it is in the presence of the drugs, the more synergistic the drugs are. This correspond to more positive ComboScore values.

Supplementary Figure 1 demonstrates the distribution of observed ComboScore values between all three screening centers. University of Pittsburg and SRI International both have a close number of data instances, about 2000 combinations per cell line. NCI’s Frederick National Laboratory, on the other hand, contains less information: about 200 combinations per cell line are tested in this center. The distribution of observed values in first two seems similar, however, SRI International reports more extreme (highly negative or highly positive) ComboScores. In total, 90% of calculated ComboScores are in range between -100 and 100, and only 0.1% are outside of the range from -500 to 500.

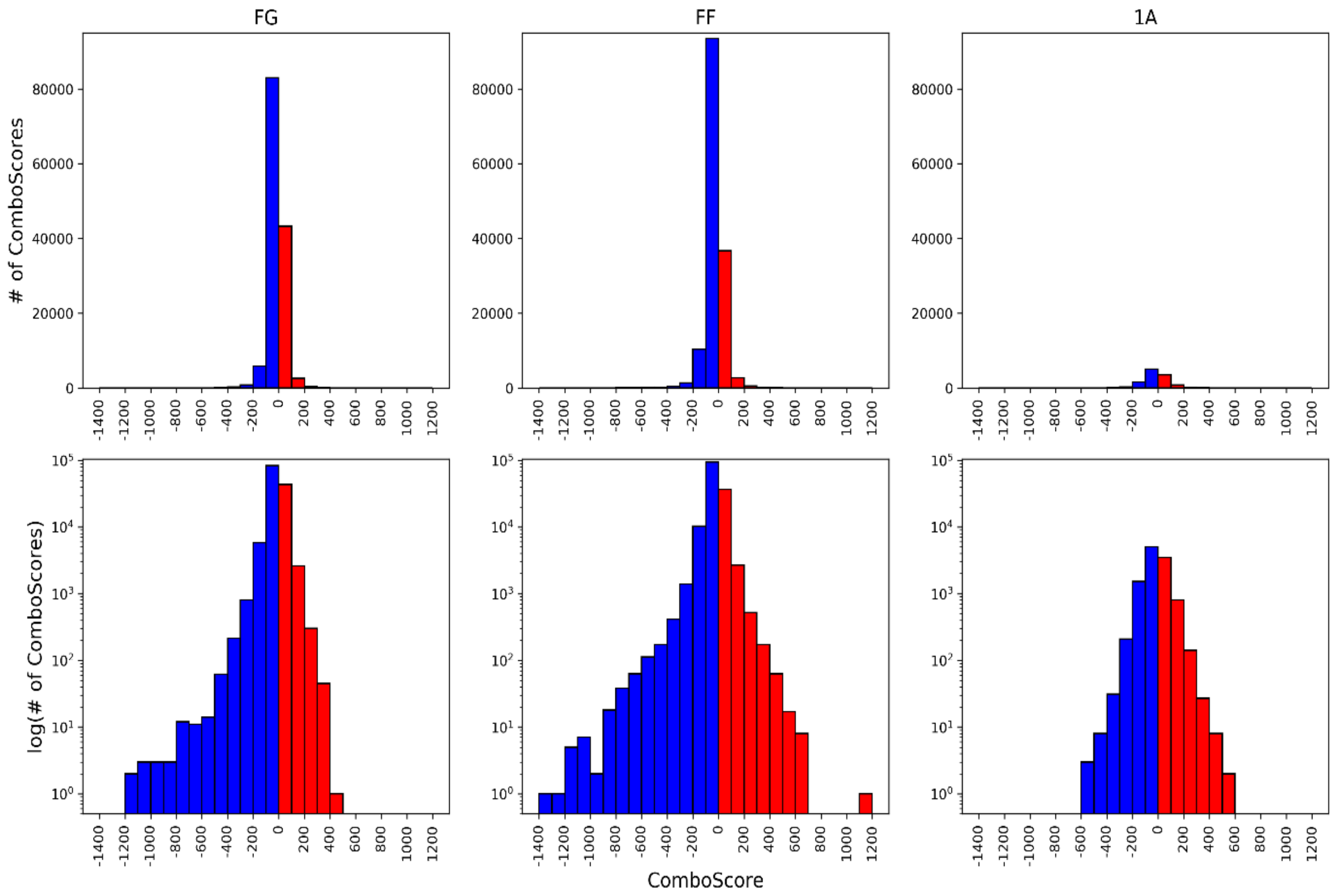

**Supplementary Figure 1.** Histograms of ComboScores measured by each screening center. Number of drug combination – cell line pairs in normal (top row) and logarithmic (bottom row) scales for antagonistic (negative, in blue) and synergistic (positive, in red) effects are presented.

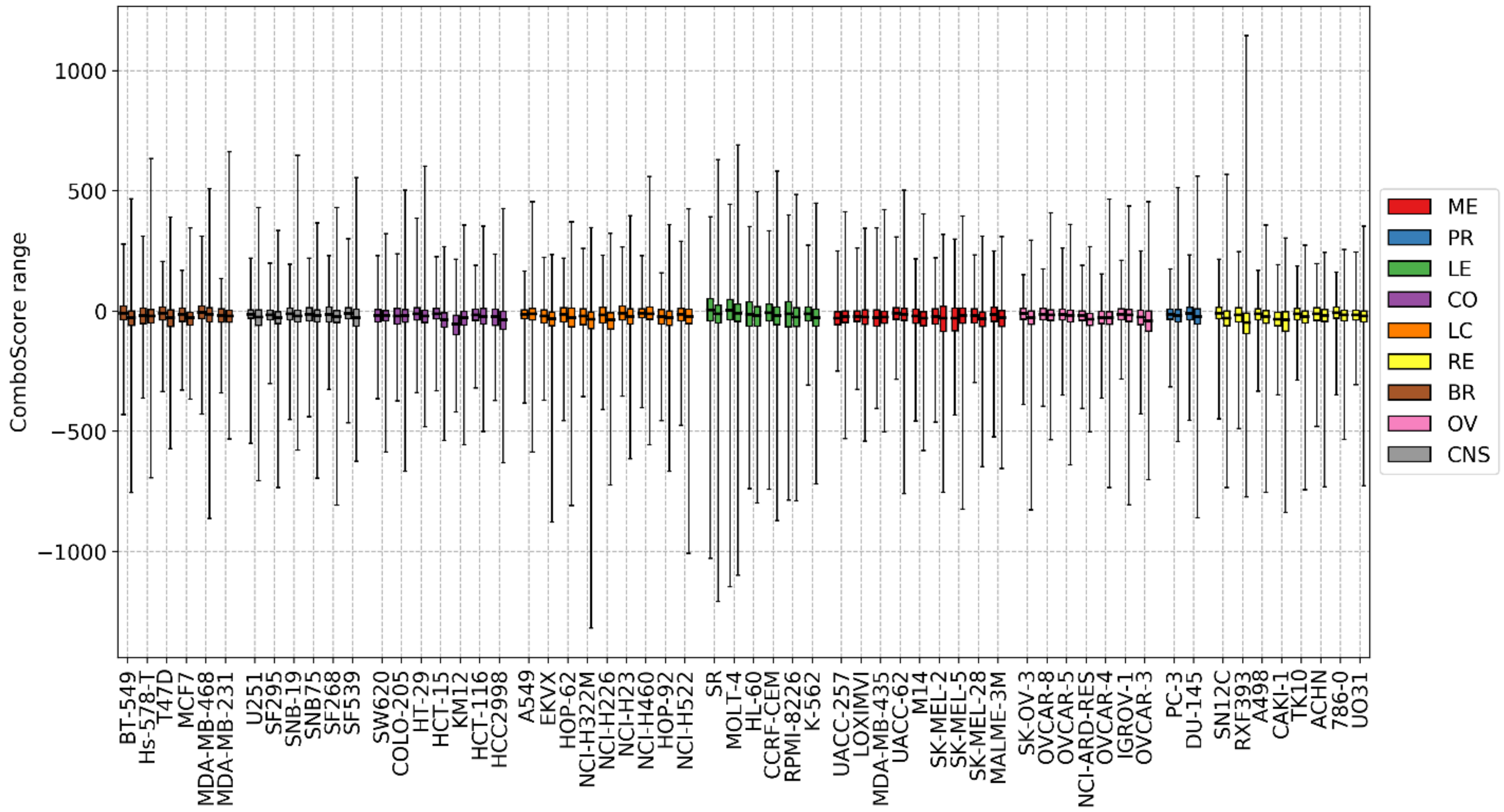

**Supplementary Figure 2.** ComboScore value ranges per cell line. For each cell line, left box corresponds to FG dataset, right box – to FF dataset. The ranges are consistently larger for FF dataset. Boxplots are colored following the cancer type.

**Types of drug features**

For the use in machine learning, the structures of compounds must be encoded as vectors of numerical features – molecular descriptors [2]. Several types of structural descriptors have been considered in this work:

1) Morgan fingerprints are topological descriptors describing the connectivity of the molecular structure, which take values 0 or 1, depending on whether the pattern is present in the molecule or not [3]. They have been calculated with RDKit library [4] using following parameters – length is 256 bits, radius is 2.

2) Morgan fingerprint counts – same as above, but instead of 0 and 1 they take integer values equal to the number of times the pattern is detected in the molecule. 256 features per drug.

3) MACCS keys encode presence or absence of 166 predetermined substructural fragments as binary vectors. Calculated with RDKit.

4) ISIDA fragments encode structure as a vector of numbers of occurrences of substructural fragments of given nature and topology in the molecule [5]. Calculated with ISIDA/Fragmentor [6]. Only one type of fragments is considered here: sequences of atoms and bonds of length 2 to 6. 1325 features per drug in total.

5) SIRMS fragments – number of occurrences of 4-atom fragments of varying topology in a molecule, including bonded and non-bonded atoms [7]. Calculated with SiRMS python library (github.com/DrrDom/sirms). 1454 features per drug.

In addition to these, 7 physico-chemical features are calculated by RDkit: total polar surface area (TPSA), molecular weight, logP, number of aliphatic and aromatic rings, H-bond donors and acceptors. They may be added to the initial pool of features.

**Predictive Performance Metrics**

To evaluate a model's performance, following parameters are calculated from observed $y_{obs}$ and predicted $y_{pred}$ ComboScore values:

1) Root Mean Squared Error (*RMSE)*:

$$RMSE = \sqrt{\frac{\sum_N \left(y_{i,obs} - y_{i,pred}\right)^2}{N}}$$

2) Coefficient of determination (*$R^2$)*[8]:

$$R^2 = 1 - \frac{\sum_N \left(y_{i,obs} - y_{i,pred}\right)^2}{\sum_N \left(y_{i,obs} - \overline{y_{obs}}\right)^2} = 1 - \frac{RMSE^2}{Var(y_{obs})}; \quad \overline{y_{obs}} = \frac{1}{N}\sum_{i=1}^{N} y_{i,obs}$$

3) Pearson's correlation coefficient ($R_p$):

$$R_p = \frac{\sum_N (y_{i,obs} - \overline{y}_{obs})(y_{i,pred} - \overline{y}_{pred})}{\sqrt{\sum_N (y_{i,obs} - \overline{y}_{obs})^2}\sqrt{\sum_N (y_{i,pred} - \overline{y}_{pred})^2}}$$

4) Spearman's rank-order correlation coefficient ($R_s$):

$$R_s = R_p(\text{rank}\, y_{obs}, \text{rank}\, y_{pred})$$

We use Pearson correlation coefficient $R_p$ between observed and predicted values of ComboScore of a dataset not used to train the model as a primary metric of its accuracy.

**Per-cell line 10-fold cross-validation on FG datasets**

Standard k-fold cross-validation proceeds as following: the dataset is randomly divided in k parts, one is left out as a test set, and other k-1 parts are used to build a model, which is then evaluated on the left-out subset. It is repeated for every subset, so that each instance of the set is predicted exactly once. 10-fold cross-validation has been performed for the RF and XGB cell line models of FG screening center data to confirm the findings of the initial validation on a similar-sized (10% of the set) test sets. All cross-validations of XGB models are carried out with the recommended values for XGBoost's hyperparameters, as comprehensively tuning in random data partitions only provided marginal gains despite the far higher computing time required.

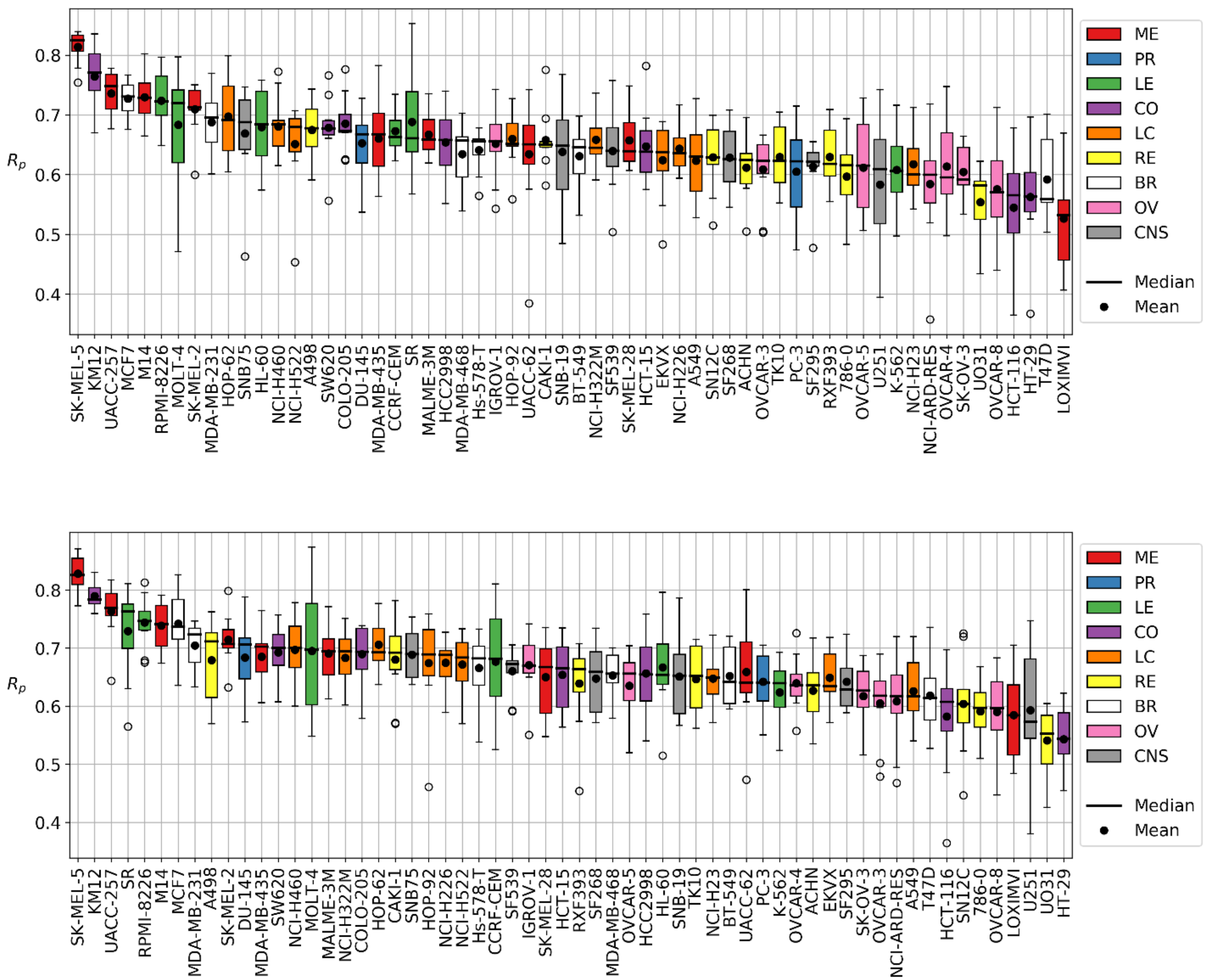


**Supplementary Figure 3.** Test set prediction performance (Pearson correlation $R_p$) in 10-fold-cross-validation with Random Forest (top) and XGB (bottom) per cell line. Both algorithms use the recommended values for their hyperparameters. Each boxplot represents the distribution of performances across test folds. Mean and median $R_p$ are indicated, boxes are sorted by median performance. Color code indicates cancer tissue type. Some tissue types (such as melanoma ME and leukemia LE) demonstrate higher overall performance than other (e.g. renal cancer RE). Correlation between 10-fold CV results (mean $R_p$ between folds) and results of random test set prediction across cell lines (as in the exploratory part) by RF is $R_p$=0.56, by XGB $R_p$=0.60.

**Per-drug Leave-One-Drug-Out cross-validation on FG datasets**

Leave-one-drug-out cross-validation is carried out in each of the 60 cell lines. Collectively, this results in a RMSE for each left-out drug and cell line pair. Rearranging these results per drug permits assessing how well the left-out drug is predicted across the 60 cell lines (Supplementary Figure 4).

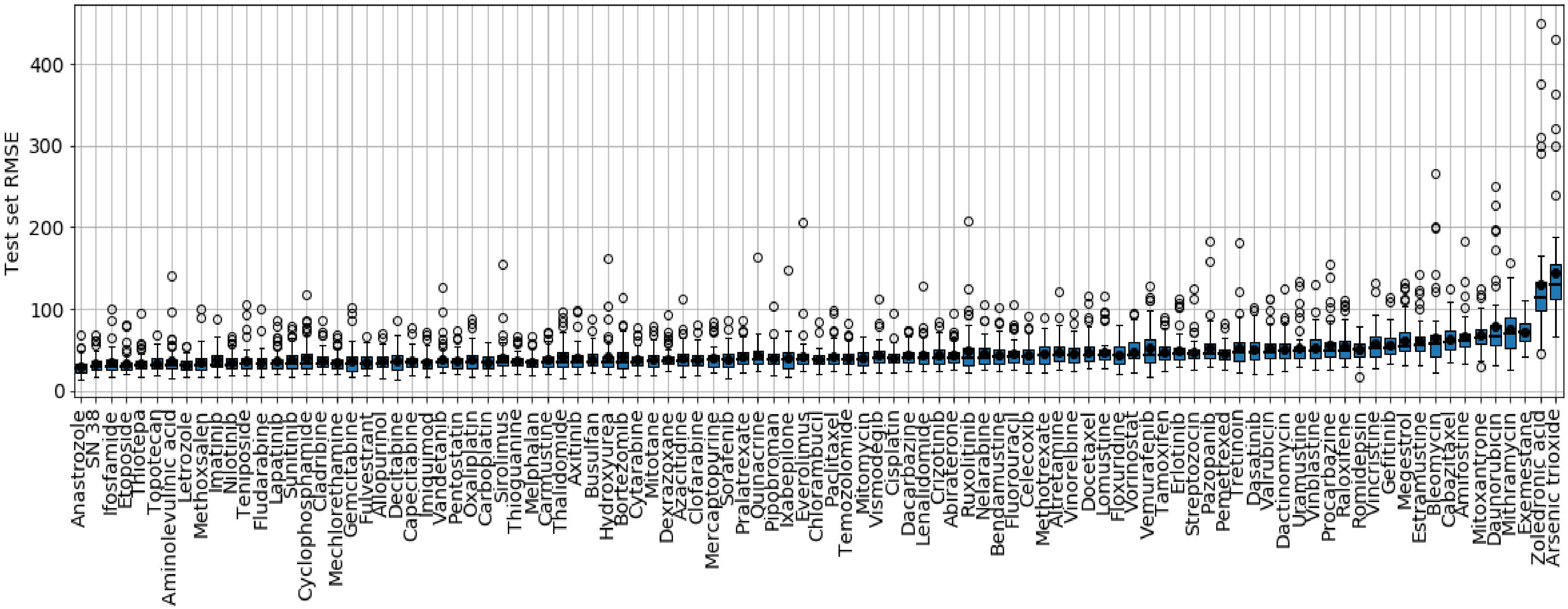
Test set RMSE
400
300
200
100
0
Anastrozole
SN 38
Ifosfamide
Etoposide
Thiotepa
Topotecan
Aminolevulinic acid
Letrozole
Methoxsalen
Imatinib
Nilotinib
Teniposide
Fludarabine
Lapatinib
Sunitinib
Cyclophosphamide
Cladribine
Mechlorethamine
Gemcitabine
Fulvestrant
Allopurinol
Decitabine
Capecitabine
Imiquimod
Vandetanib
Pentostatin
Oxaliplatin
Carboplatin
Sirolimus
Thioguanine
Melphalan
Carmustine
Thalidomide
Axitinib
Busulfan
Hydroxyurea
Bortezomib
Cytarabine
Mitotane
Dexrazoxane
Azacitidine
Clofarabine
Mercaptopurine
Sorafenib
Pralatrexate
Quinacrine
Pipobroman
Ixabepilone
Everolimus
Chlorambucil
Paclitaxel
Temozolomide
Mitomycin
Vismodegib
Cisplatin
Dacarbazine
Lenalidomide
Crizotinib
Abiraterone
Ruxolitinib
Nelarabine
Bendamustine
Fluorouracil
Celecoxib
Methotrexate
Altretamine
Vinorelbine
Docetaxel
Lomustine
Floxuridine
Vorinostat
Vemurafenib
Tamoxifen
Erlotinib
Streptozocin
Pazopanib
Pemetrexed
Tretinoin
Dasatinib
Valrubicin
Dactinomycin
Uramustine
Vinblastine
Procarbazine
Raloxifene
Romidepsin
Vincristine
Gefitinib
Megestrol
Estramustine
Bleomycin
Cabazitaxel
Amifostine
Mitoxantrone
Daunorubicin
Mithramycin
Exemestane
Zoledronic acid
Arsenic trioxide

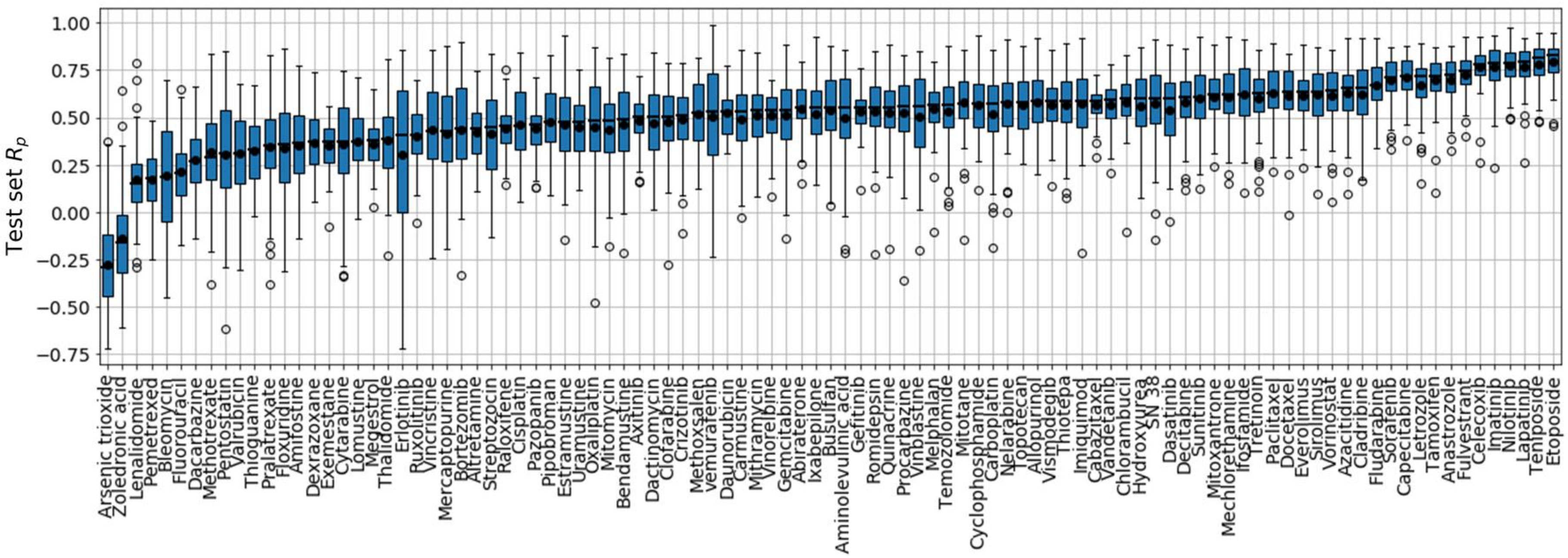


**Supplementary Figure 4**. **(top)** Test set prediction performance (RMSE) in leave-one-drug-out cross-validation with XGB (using the recommended values for their hyperparameters) per left-out drug. Each boxplot represents the distribution of scores across cell lines. Mean and median scores are indicated. The boxplots are sorted in order of median RMSE, from lowest to highest. The distribution of predictions is similar to previously discussed RF models: models for Arsenic trioxide and Zoledronic acid have the highest median prediction error (>100), Anastrazole and SN 38 are again in the lead. **(bottom)** Test set prediction performance ($R_p$) using the same set of predictions as above. Models for Arsenic trioxide and Zoledronic acid have the lowest median prediction performance ($R_p<0$), whereas tyrosine kinase inhibitors (Imatinib, Nilotinib, Lapatinib) and topoisomerase inhibitors (Teniposide, Etoposide) are among the best-predicted left-out drugs.

**Per-cell line Leave-One-Drug-Out cross-validation on FF datasets**

We subjected the FF dataset to the same LODO cross-validation analysis as FG. Supplementary Figure 6 shows the results of LODO cross-validation for FF screening center data. The results of the validation for FF datasets are substantially worse than for FG across all cell lines. Only the top 25% models have average performance $R_p$>0.35 (Supp Figure 6). Therefore, we may conclude that there are some inconsistencies within the FF dataset. The most notable occurrence here is the abundance of models obtaining $R_p$=1 or -1 across all cell lines. These models correspond to left-out drugs that were only tested with two partners. While this does never occur in the FG dataset, 20 of the 92 drugs tested in the FF are only partnered with doxorubicin and triethylenemelamine (both drugs are absent from FG). Thus, for these drugs the correlation coefficient will be 1 or -1, depending on whether the models are able to correctly learn the synergy trend of these two left-out ComboScores.

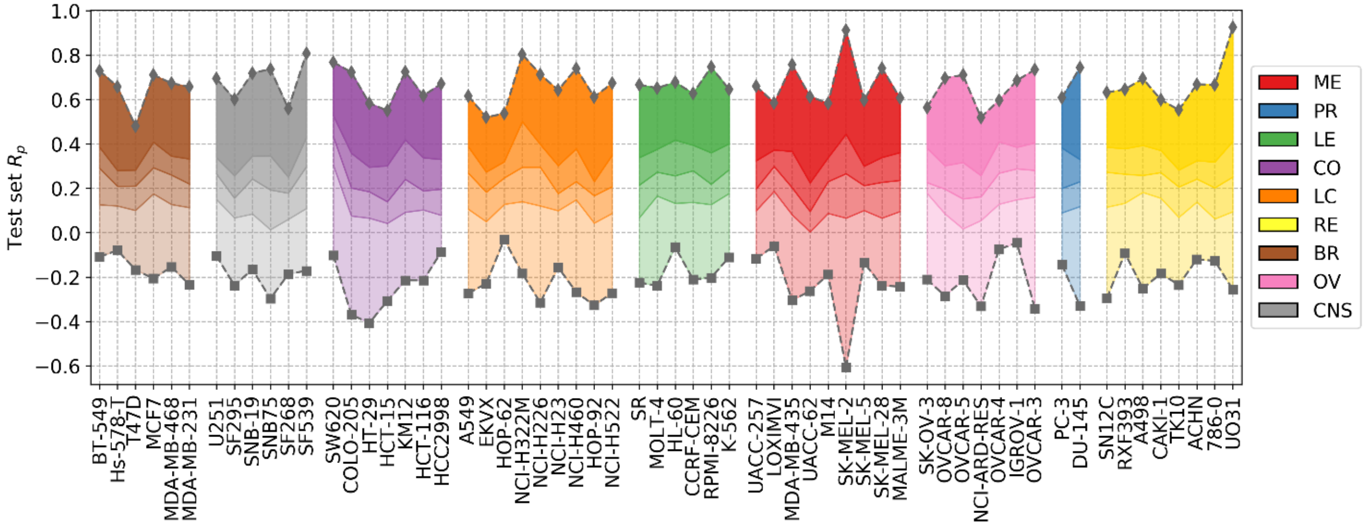


**Supplementary Figure 5.** Leave-one-drug-out cross-validation results with XGB (using the recommended values for their hyperparameters) trained on FF datasets, when using only drugs that have 3 or more partners (i.e. three or more test set instances). Distribution of models' performances is shown by cancer type (color code). Each colored zone represents 25% of models per cell line: from dense zone – top performing 25%; to light zone – bottom quartile. The performance is not changed compared to the one using all data (Figure 7 in main text), with median $R_p$ across cell lines ranging from 0.214 (average for prostate cancer PR, in blue) to 0.254 (average for leukemia LE, in green). Maximum performances range from 0.643 (average for ovarian cancer OV, in pink) to 0.686 (average for brain cancer CNS, in grey) on average across cell lines. Minimum performances range from -0.243 (average for colorectal cancer CO, in purple) to -0.159 (average for breast cancer BR, in brown) on average across cell lines.

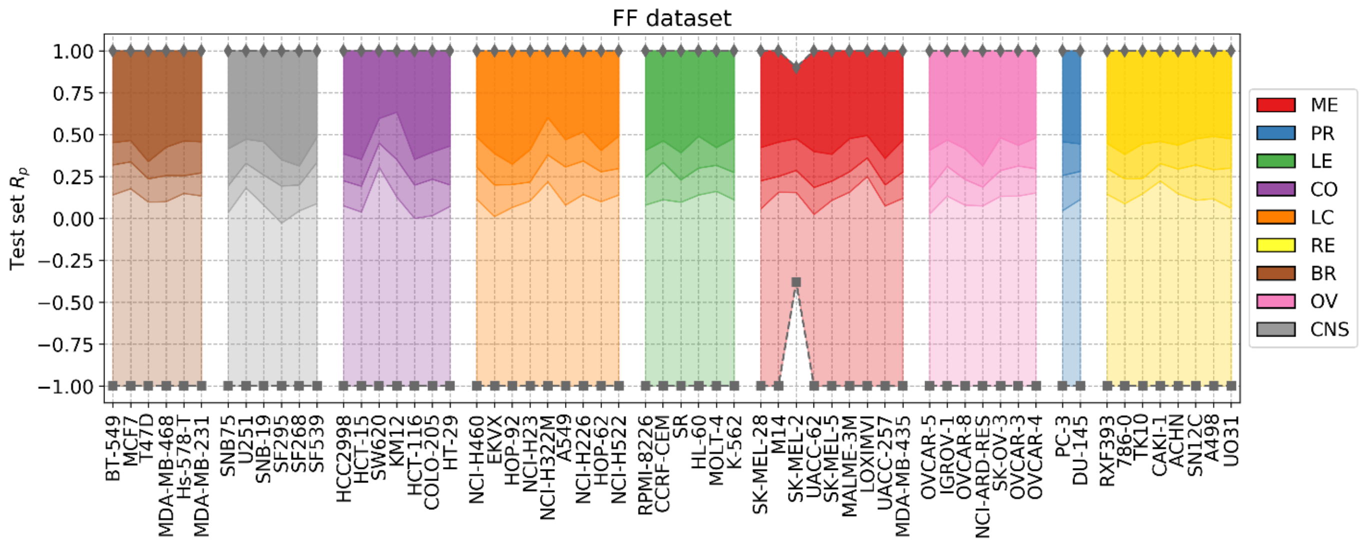


**Supplementary Figure 6.** LODO cross-validation results with XGB (using the recommended values for their hyperparameters) on the FF datasets. Distribution of models' performances is shown by cancer type (color code). Each colored zone represents 25% of models per cell line: from dense zone – top performing 25%; to light zone – bottom quartile. The method demonstrates lower accuracy on this dataset, with median $R_p$ across cell lines ranging from 0.214 (prostate cancer PR, in blue) to 0.254 (leukemia LE, in green), compared to the range from 0.479 (RE) to 0.554 (ME) for FG datasets (see Figure 6). Performances equal to 1 and -1 correspond to drugs for which only two partners are available, thus, only two combinations for that left-out drug are present in the test set of that cell line.

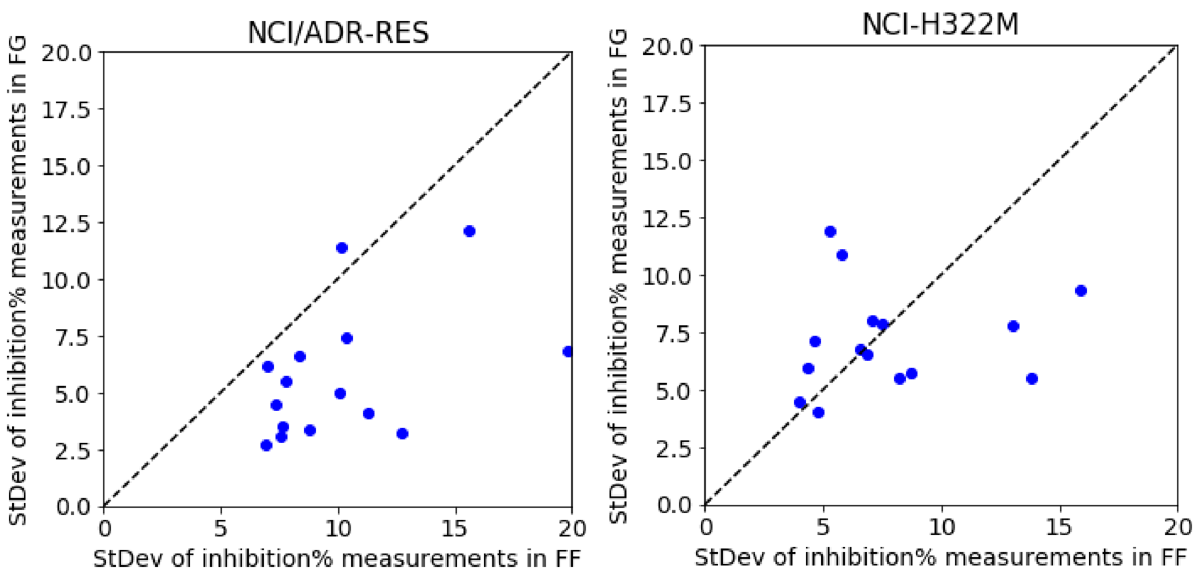


**Supplementary Figure 7.** Random Forest (RF) model performance comparison for FG and FF datasets. Models are built following the final setup in the exploratory analysis (*i.e.* a 90/10 data partition is employed for each cell line; MFPC with physico-chemical features as well as data augmentation are also used). We analyzed five drugs (Thioguanine, Chlorambucil, Altretamine, Fluorouracil and Melphalan) with a high number of different test dates in both centers. On the left, the results with the cell line that is worst predicted by RF with FF data (NCI/ADR-RES with Rp=0.14 in 90/10 partition), which is much better predicted with FG (Rp=0.65, using the same partition). This plot shows the standard deviation of the values of each set from FG against those from FF. 14 of the 15 sets have higher standard deviation of PERCENTGROWTH with FF data, which is consistent with the low predictive performance obtained with this dataset. On the right, we repeated this operation with the cell line that is better predicted by RF with FF data (NCI-

H322M with Rp=0.61 in 90/10 partition), which is also well predicted with FG (Rp=0.66, always with the same partition). By contrast, this plot shows that only 7 of the 15 sets have higher standard deviation with FF data (the same five drugs are used).
